# Cross Domain Consistency of Aesthetic Preference-driven Social Behavior

**DOI:** 10.64898/2026.03.21.713367

**Authors:** Trung Quang Pham, Junichi Chikazoe

## Abstract

Aesthetic preference is a primary driver of social behavior in the digital era, yet the extent to which these preferences remain consistent across disparate domains remains poorly understood. We hypothesize that aesthetic judgment is governed by a domain-invariant latent structure, such that individuals who exhibit similar preferences in one category will demonstrate comparable alignment in seemingly unrelated domains. To test this, we recruited 37 participants to evaluate stimuli across three distinct aesthetic domains: art, faces (male and female), and scenes. We developed a novel computational framework that reformulates cross-domain preference as a user-based collaborative filtering problem, encoding individual profiles through inter-subject similarity matrices. Our model successfully predicted participant responses in a target domain based on their similarity to the cohort in a separate source domain. These results demonstrate robust cross-domain consistency, suggesting that aesthetic evaluation is mediated by an abstract, domain-general mechanism rather than being purely stimulus-dependent. We propose that this consistency is rooted in a shared neurophysiological pathway, likely involving the orbitofrontal cortex (OFC) and the Default Mode Network (DMN), and discuss how these findings provide a foundation for more sophisticated, cross-modal recommendation systems and the study of individual social identity.

## 1 INTRODUCTION

Aesthetic preference is a fundamental factor that drives our social behavior in the digital age. Research indicates that visual content in an online review appears to attract higher engagement than the text itself (Li et al. (2024)), highlighting a significant shared preference for attractiveness. While aesthetic preference has long been considered highly variable across cultures (Masuda et al. (2008); Chen et al. (2016); Yang et al. (2019)), domains (Vessel et al. (2018)), and modalities (Lee et al. (2025)), recent behaviors and cognitive evidence suggests a dual nature of aesthetic preferences, characterized by both idosyncratic specificity and underlying universality.

Miller and Hübner (2020) proposed a Theory of Aesthetic Preferences (TAP), which has since been extended to demonstrate cross-cultural universality (Chinese-German; Miller et al. (2025); Japanese and German; Mikuni et al. (2024)). TAP suggests that aesthetic preferences are driven more by the intrinsic properties of the stimuli than by culture, with universal aesthetic responses being closely linked to positive emotions. While this theory effectively explains how people infer the aesthetic preferences of others both within and across different cultures, it raises a fundamental question: are these preferences consistent across different domains?. To date, empirical work has focused almost exclusively on the “art” domain (Miller and Hübner (2020); Iigaya et al. (2021, 2023); Miller et al. (2025)), leaving the generalization to other domains largely unexplored.

From a neurophysiological perspective, functional Magnetic Resonance Imaging (fMRI) studies have identified the orbitofrontal cortex (OFC) and the lateral prefrontal cortex (lPFC) as central hubs for aesthetic preferences (Jacobsen et al. (2006); Ishizu and Zeki (2013); Ticini (2017); Iigaya et al. (2023)). The recruitment of these areas appears to depend on whether the aesthetic appraisal is action-oriented. If the aesthetic appraisal occurred during the simulation of an experience, the lPFC is typically engaged. In contrast, post-stimulus appraisal activates the OFC more frequently. Given that both OFC and lPFC are high-level integration areas known for domain-independent processing(Chikazoe et al. (2014)), these findings suggest that aesthetic preferences may be encoded as abstract, domain-general information.

In the present study, we evaluate the cross-domain consistency of aesthetic preference using a novel computational framework. Three distinct aesthetic domains were selected, i.e., art, face (male and female), and scene, to test our hypothesis. By proposing an aesthetic preference encoding approach based on cross-subject similarity matrices, we developed a cross-domain behavioral model designed to predict individual responses across different categories. Our results demonstrate that a participant’s preferences in one domain can accurately predict their behavior in another. Consequently, monitoring individual aesthetic preferences provides valuable insights for the study of modern social behavior and the optimization of digital recommendation systems.

## 2 HYPOTHESIS & MODELING APPROACH

We hypothesize that while the behavioral responses may vary across contexts, the underlying latent preference structure remains invariant across domains and serves as the primary driver of individual behaviors. If this hypothesis holds, an individual’s preference profile identified in one domain should possess the predictive power to forecast their response in a distinct domain (Fig. 1A).

**Figure 1.**
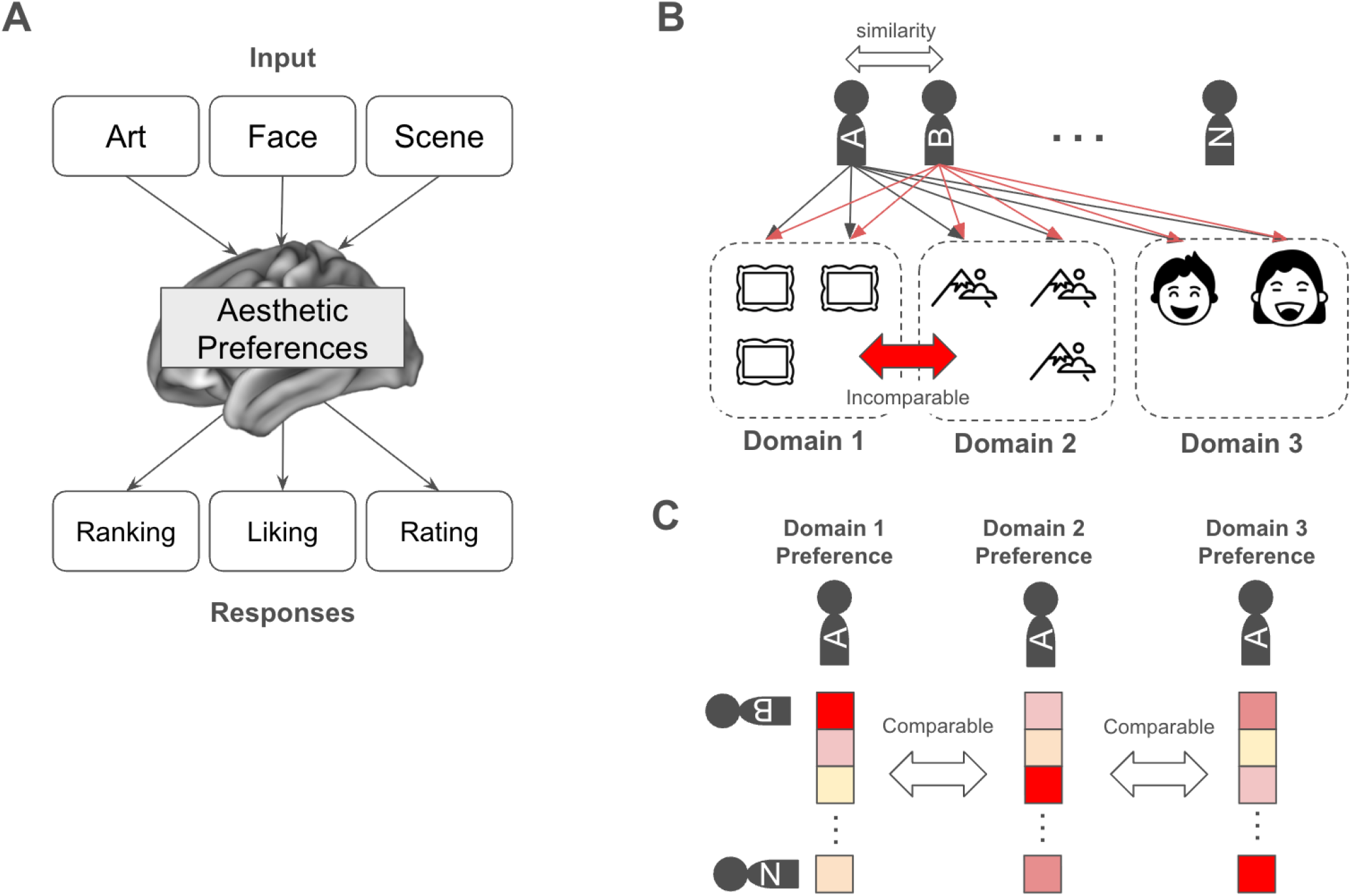
Conceptual framework of aesthetic-driven social behavior across domains. (A) Schematic representation of heterogeneous inputs and behavioral responses across the three selected domains. (B) The mapping of individual aesthetic preferences onto a collective space, modeled as a user-based collaborative filtering problem to bypass intrinsic variance in domain-specific features. (C) Encoding of individual preference profiles through an inter-subject similarity matrix.

Representing the personal preferences across disparate domains presents a significant computational challenge. Traditional aesthetic modeling has heavily relied on rule-based systems and expert-derived criteria (O’Donovan et al. (2014)). More recently, research has utilized artificial neural networks to extract generic, though often “black-box” or uninterpretable, image features (Murray et al. (2012); Lu et al. (2015); Brachmann and Redies (2017); Iigaya et al. (2021)). However, due to the inherent differences in features and scaling across domains, direct within-participant response mapping is less reliable.

To address this, we propose that if preferences are consistent across domains, the relative relationship between a participant and the broader population should also remain stable. By anchoring an individual’s responses to those of others via inter-subject similarity, we can derive a simpler and more robust representation of aesthetic preference (Fig. 1B). This approach is predicated on the condition that the participant cohort is rigorously controlled, specifically, i.e., the same group of individuals is maintained across all experimental domains.

This framework reformulates individual social behavior as a typical user-based collaborative filtering task, where similarity vectors represent a participant’s aesthetic standing within a population or society (Fig. 1C). Consequently, participant behavior can be predicted using a standardized user-based collaborative filtering formula, defined as follows,

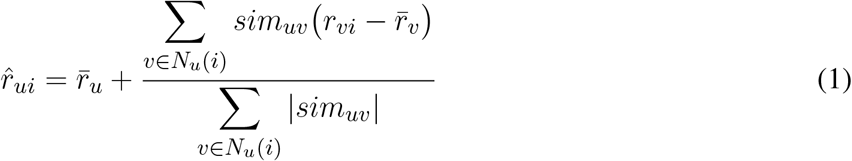

where:

- 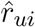: predicted response of participant *u* for item *i*
- 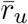: average response of participant *u*
- *N*_*u*_(*i*): neighborhood of users similar to *u* who rated item *i*
- *sim*_*uv*_: similarity between participant *u* and participant *v*
- *r*_*vi*_: response of participant *v* for item *i*

## 3 MATERIALS AND METHODS

### 3.1 Experiments

A total of 37 participants (9 males, 28 females, all Japanese nationality) were recruited for behavioral experiments, in which they evaluated items across three distinct aesthetic domains: art, faces (male and female), and scenes. The gender imbalance was the result of a non-stratified sampling during the recruitment phase and was not intentionally specified.

The stimulus set is comprised of 400 items for art, 160 items for faces (80 male and 80 female), and 400 items for scenes. The scene images were randomly sampled from the Aesthetic Visual Analysis dataset (AVA; Murray et al. (2012)), a standard benchmark for aesthetic assessment. The art images were downloaded from LiveAuctioneers (https://www.liveauctioneers.com/). The face images were collected exclusively from printed publications, such as commercial magazines, and subsequently digitized; recognizable celebrities were excluded to prevent the influence of prior personal associations. The variation in sample sizes (400 vs. 160) was dictated by practical constraints during the stimuli preparation phase and was not intended to reflect a hierarchy of importance between domains. To ensure the independence of the evaluations, there was no overlap of stimulus items between the three domains.

The experimental procedure consisted of two distinct phases: a subjective valuation task and an ordinal ranking task. In the first phase, participants were asked to provide subjective monetary valuations for artworks (Fig. 2A). The metric was chosen to capture the high-resolution, information-rich measurement of aesthetic preference. The target artwork was displayed in the upper panel, with the associated price shown below. Participants adjusted the price using up/down arrow keys, starting from a baseline of JPY 10,000, within a permissible range of JPY 100 to JPY 10,000,000.

**Figure 2.**
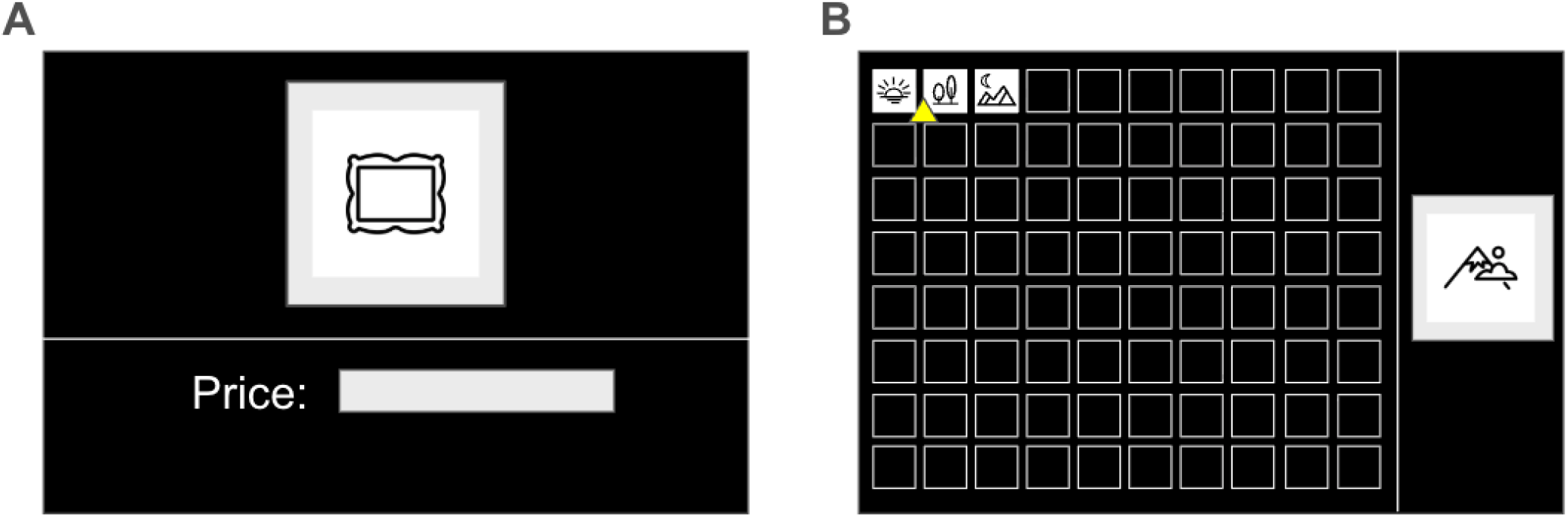
Illustrations of experimental User Interface. (A) Interface for artwork rating task. Artworks were evaluated individually, with a default starting value of JPY 10,000. (B) Interface for the facial and scene ranking tasks. The yellow triangle highlights the currently selected position within the 8 ×10 grid. Ranking follows a left-to-right, top-to-bottom progression, where the top-left position presents rank 1 and the bottom-right presents rank 80.

In the second phase, participants were asked to rank the human faces and scenes (Fig. 2B). This transition in methodology was necessitated by the fact that human faces do not lend themselves to appropriate monetary valuation. During this task, stimuli were presented sequentially in a right-side panel, while previously ranked items were organized in an 8 × 10 grid on the left. Participants navigated the grid using keyboard arrow keys to assign a relative rank to the current stimulus, confirming the selection with the “Enter” key. Ranking values ranged from 1 to 80. For the scene domain, the 400 items were randomly pre-divided into five subsets of 80 items each, with each subset ranked independently to maintain consistency with the grid format.

All stimuli were presented on a 29-inch LCD monitor, using Presentation® software (v23.0, Neurobehavioral Systems, Inc., Berkeley, CA, www.neurobs.com). To mitigate potential order effects, the presentation sequence of both the items and the domains was fully randomized across all participants. All the ranking and rating behaviors reflect individual value judgments that may incorporate their socially learned norms.

### 3.2 Data-processing

Since the experimental data were collected across different time frames and under varying constraints, the raw response distributions exhibited significant variance. To ensure comparability across domains, we applied z-score normalization to the participant responses within each domain. The normalized values, *x*_*norm*_ were calculated as follows:

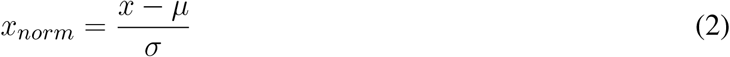

where *x* is observed raw response, *µ* is the domain-specific mean, and *σ* is the corresponding standard deviation. To prevent data leakage and maintain consistency across all analyses, the dataset for each domain was partitioned into training and testing subsets using a randomized 75/25 split (3:1 ratio). This partitioning was performed prior to any modeling or feature extraction to ensure the integrity of the cross-domain validation.

### 3.3 Similarity analysis

The preference similarity between two participants, *u* and *v*, in a single domain, *sim*_*uv*_, is measured by the Pearson correlation as follows:

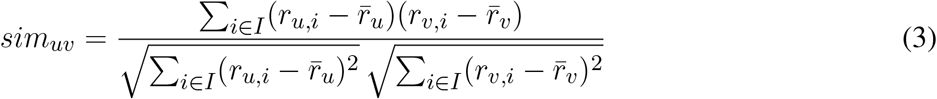

where *r*_*u,i*_ is participant *u*’s response to item *i*, and *r*_*v,i*_ is participant *v*’s response to item *i*. This procedure results in a 37 × 37 similarity matrix for each domain.

To calculate pattern similarity across domains, we vectorized the lower triangle of each domain’s similarity matrix and then calculated the Pearson correlation coefficients between these vectors, resulting in a 4 × 4 correlation matrix.

### 3.4 Collaborative filtering model

There are many available methods for solving the collaborative filtering task. Among them, the Sparse Linear Method (SLIM) and its variants consistently demonstrate outstanding performance. For the sake of simplicity, SLIM is adopted as in Ning and Karypis (2011), where the objective is to learn W as the minimizer of the following regularized optimization problem:

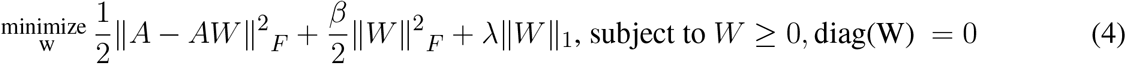

*W* is a sparse *n* × *n* matrix of aggregation coefficients between participants, which is inversely equivalent to *sim*_*uv*_ in Eq. 1. The norm ∥ · ∥ _*F*_ denotes the matrix Frobenius norm, and ∥ *W*∥ _1_ denotes the entry-wise *𝓁*_1_-norm of *W* . *A* is the item–participant rating matrix of size *m × n*, with *m* items and *n* participants. *β* and *λ* are regularization parameters. Equation 4 was implemented using the ElasticNet model from the scikit-learn library (Pedregosa et al. (2011)) in the Python environment (v3.10).

To find the optimal parameters, we performed a grid search with

- *β* in the range [10^−4^, 10^12^]
- *λ* in the range [10^−3^, 10^2^]

For each pair of *β* and *λ*, we randomly divided the training set into two halves (one for model fitting and one for validation). The *β*–*λ* pair associated with the highest performance on the validation set was selected for the final training on the full training set.

### 3.5 Evaluation

The model is evaluated by calculating the mean squared error (MSE) and the correlation between the predicted responses and the true responses of the validation set. To avoid data-splitting bias, the validation process was repeated 20 times for each pair of parameters. The average score is then used for the final parameter selection of the model.

The MSE formula is given as follows:

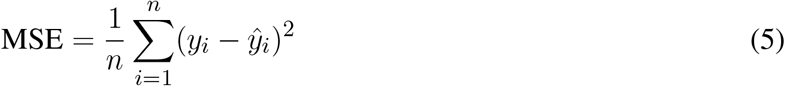

where *i* is the item index, *n* is the total number of responses, *y*_*i*_ is the observed raw response, and 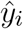 is the model-predicted response.

The similarity between the predicted responses and the observed raw responses is calculated using Pearson correlation (Eq. 3) for each participant. The correlation coefficients are transformed into Fisher’s z-scores, averaged, and then transformed back to obtain the group similarity. The Fisher z-score is computed as follows:

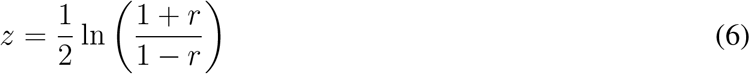

where *r* is the observed Pearson correlation coefficient.

### 3.6 Statistical analysis

Statistical analyses were conducted using Python v3.10 with extensive use of the numpy, scipy, and scikit-learn libraries. All training and post-analysis were performed on a workstation with the following specifications: Intel Core i7 CPU, 128 GB RAM, Quadro P6000 GPUs, running Ubuntu 16.04 LTS.

Statistical comparisons between Pearson’s correlation coefficients for two distinct conditions were conducted using a two-tailed paired t-test after Fisher’s z transformation (Eq. 6). The paired t-test formula is given as follows:

Where

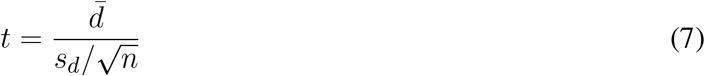

- 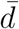 is the sample mean of the differences,
- *s*_*d*_ is the sample standard deviation of the differences,
- *n* is the number of paired observations.

Explicitly,

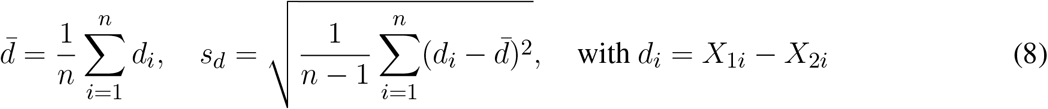

where *X*_1*i*_ and *X*_2*i*_ are the *i*th items in the first and second arrays being compared, respectively.

## 4 RESULTS

### 4.1 Behavioral analysis

We first characterized the aesthetic preferences of the participants across all domains by evaluating the similarity between their respective correlation matrices (Fig. 3). As illustrated in Fig. 3A, the inter-subject correlation space exhibited a heterogeneous distribution, featuring both highly dissimilar pairs (indicated by lighter columns) and distinct clusters of high similarity (darker clusters) that persisted across domains.

**Figure 3.**
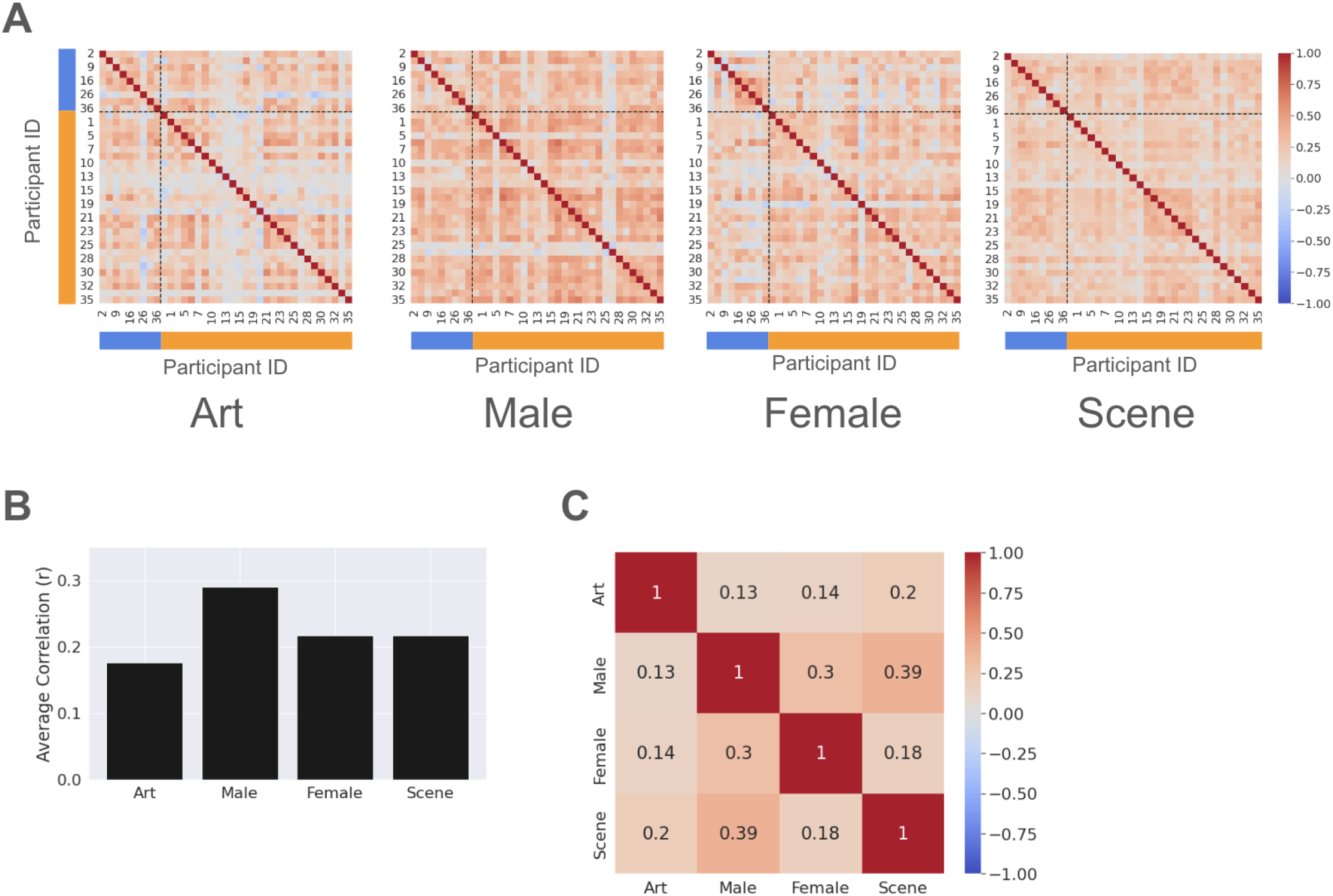
(A) Similarity matrices calculated based on: art, face (male and female), and scene datasets, respectively. Participants are reordered by gender (Males first, followed by Females) while retaining their original IDs. Axis color bars and dashed divider lines indicate the boundary between the male (blue) and female (orange) cohorts, revealing distinct blocks of intra-gender and inter-gender correlation.(B) Average similarity between participants across all domains. (C) Correlation between similarity matrices of different domains.

The mean correlation coefficient for the lower triangle matrix indicated a weak-to-moderate degree of inter-subject similarity (Fig. 3B). Notably, the highest mean value was observed within the male face stimuli set (average Pearson’s *r* = 0.289). Given that females are the dominant gender in our sample, this result suggests a potential gender specific modulation of aesthetic preferences. We further addressed such a phenomenon in subsequent sections.

Further analysis revealed a weak-to-moderate positive correlation between domain pairs (Fig. 3C). The strongest association was observed between Male and Scene sets (Pearson’s *r* = 0.39), supporting the hypothesis of cross-domain consistency in aesthetic preference. These behavioral results validate our methodological approach of utilizing cross-participant similarity as a robust anchor for representing individual aesthetic profiles.

### 4.2 Face-to-Art preferences

We subsequently evaluated the predictive power of aesthetic preferences identified in the Face domain (comprising both Male and Female subsets) to forecast responses in the Art domain. To establish a performance baseline, this cross-domain condition was compared against a within-domain “ground truth” using a leave-one-out training set.

Figure 4A shows the comparison of MSE between conditions. As expected, the MSE of the Face-to-Art condition was higher than that of the Art-to-Art control. Figure 4B shows the individual participant correlations across both conditions; notably, the correlations remained comparably high for most participants (average Pearson’s *r* = 0.55 for the Art-to-Art condition, and *r* = 0.35 for the Face-to-Art condition).

**Figure 4.**
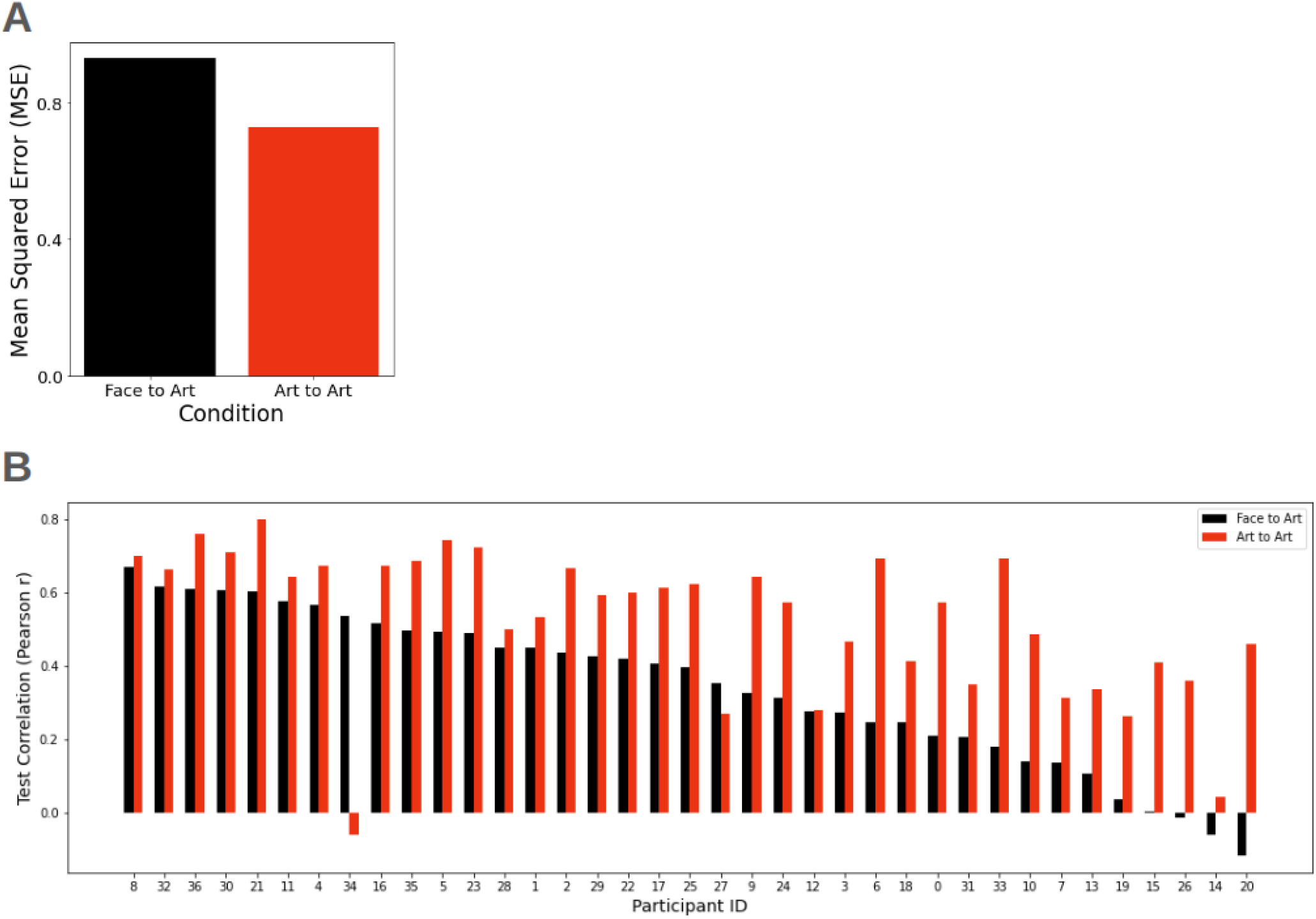
(A) Comparison of average MSE between Face-to-Art and Art-to-Art conditions. (B) Comparison of Pearson’s *r* across participants. The participant IDs are sorted in descending order of their Face-to-Art Pearson’s *r*, along the X-axis to highlight the distribution of model performance.

Similar patterns were observed in the reciprocal conditions: Face-to-Face and Art-to-Face conditions (Fig. 5). While the MSE was comparable between these two conditions, the test correlations exhibited slight variance, yielding a mean Pearson’s *r* = 0.40 for the Art-to-Face condition, compared to *r* = 0.48 for the Face-to-Face condition.

**Figure 5.**
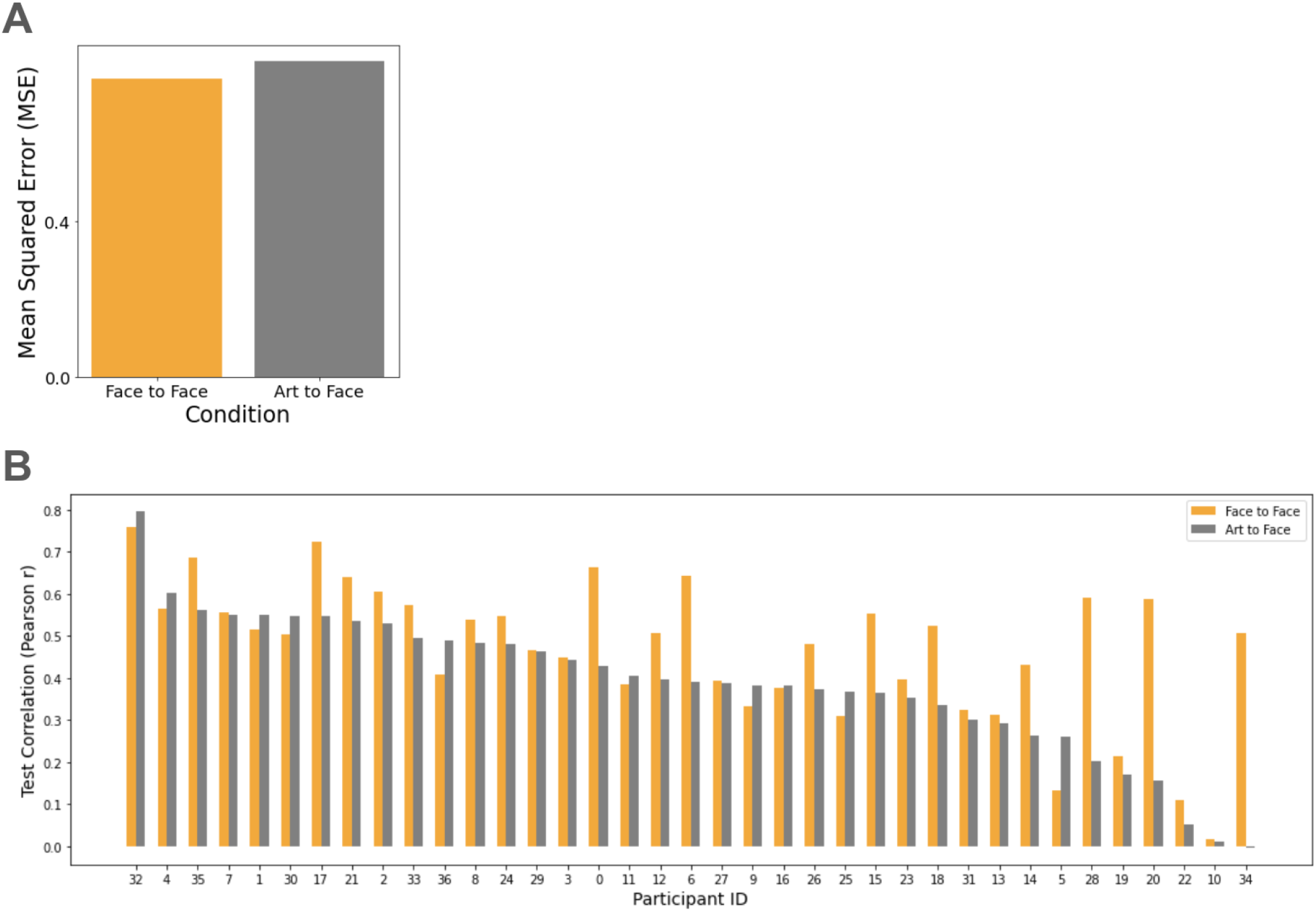
(A) Comparison of average MSE between Face-to-Face and Art-to-Face conditions. (B) Comparison of Pearson’s *r* across participants. The participant IDs are sorted in descending order of their Art-to-Face Pearson’s *r*, along the X-axis to highlight the distribution of model performance.

Taken together, these results empirically demonstrate the feasibility of cross-domain aesthetic inference. The relatively high correlation coefficients achieved in the cross-domain tests suggest that a significant portion of aesthetic preference is represented by a domain-general latent structure that can be successfully captured by our similarity-based framework.

### 4.3 Gender-specific modulation on aesthetic preferences

Next, we investigated the gender specific modulations on aesthetic preference within a cross-modal framework. We compared prediction performance between two sub-conditions of the Face-to-Art task: same-gender condition and the opposite-gender condition (Table 1). In the same-gender condition, the model was trained using participant responses to faces of their own gender (i.e., male participant responses to male faces and vice versa). Whereas, the opposite-gender condition used responses to faces of the participant’s preferred gender. Both models were then tested on art and scene prediction tasks. Due to sample size imbalance between male (*n* = 9) and female (*n* = 28) participants, we stratified the cohort by gender and evaluated each group independently.

**Table 1.** Test conditions for investigating gender influences on aesthetic preferences.

| Condition | Participant Group | Train set | Test set |
| --- | --- | --- | --- |
| Same gender | Male | Male | Art/Scene |
|  | Female | Female | Art/Scene |
| Opposite gender | Male | Female | Art/Scene |
|  | Female | Male | Art/Scene |

Figure 6 summarizes the model performance for each condition using MSE. For the male group, a modest increase in MSE (1.5 %) was observed between same-gender and opposite-gender conditions in art prediction (Fig. 6A left; paired t-test, *t*(8) = 0.46, *p* = 0.65; not significant). A greater increase in MSE (4.69 %) was found in scene prediction (Fig. 6B left; paired t-test, *t*(8) = 2.84, *p* = 0.021; statistically significant). Conversely, no significant difference was found between same-gender and opposite-gender conditions for female group in both art prediction (Fig. 6A right; paired t-test, *t*(27) = 1.59, *p* = 0.12) and scene prediction (Fig. 6B right; paired t-test, *t*(27) = 1.38, *p* = 0.17).

**Figure 6.**
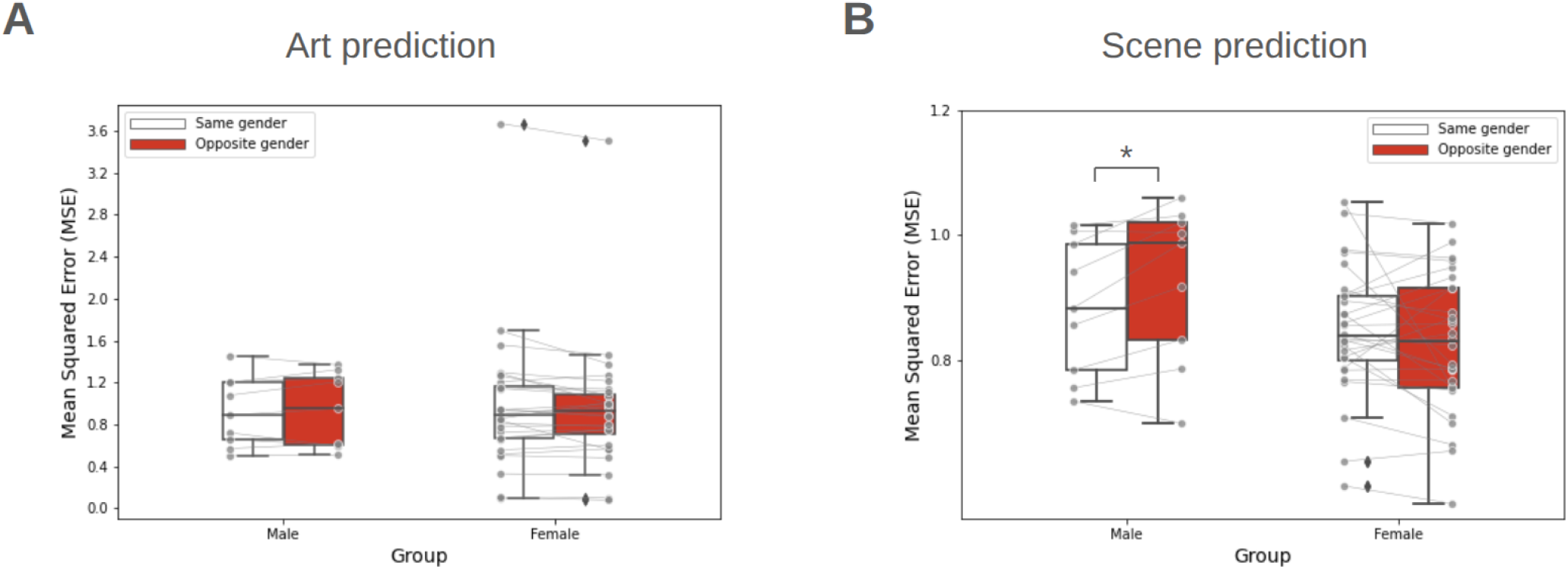
(A) Comparison of average MSE between same-gender and opposite-gender conditions in art prediction. (B) Comparison of MSE between same-gender and opposite-gender conditions in scene prediction. Center lines indicate median, box boundaries indicate inter-quartile range, and whiskers indicate the range of the remaining data, excluding outliers. ∗ : *p <* 0.05.

These findings were corroborated by comparing the average correlation coefficients between conditions (Fig. 7). For the male group, the average correlation coefficient was reduced 19.3 % in scene prediction when moving from same-gender to opposite-gender predictors (Fig. 7B left; paired t-test, *t*(8) = 2.50, *p* = 0.018; statistically significant). No other significant difference was found across the remaining cases.

**Figure 7.**
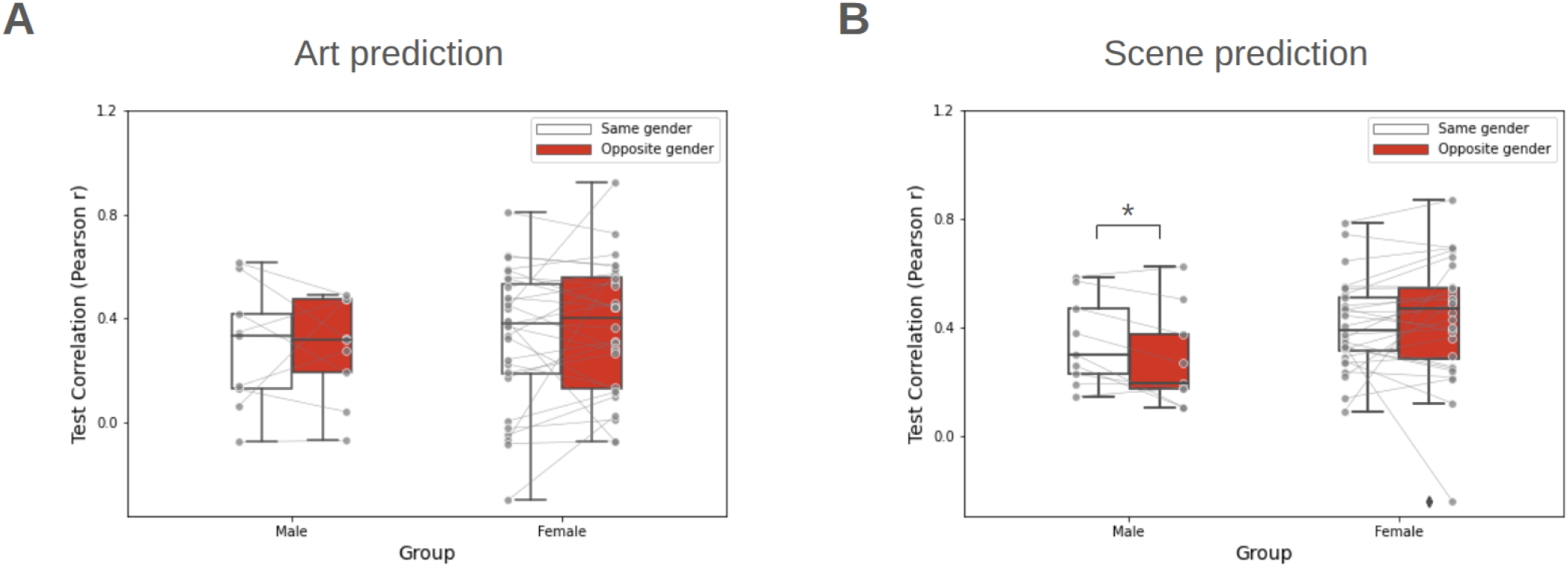
(A) Comparison of average Pearson’s *r* between same-gender and opposite-gender conditions in art prediction. (B) Comparison of Pearson’s *r* between same-gender and opposite-gender conditions in scene prediction. Center lines indicate median, box boundaries indicate inter-quartile range, and whiskers indicate the range of the remaining data, excluding outliers. ∗ : *p <* 0.05.

While the median performance metrics for the female group appeared slightly higher than those for the male group in both art and scene prediction, this was likely an artifact of the disparate sample size (*n*_*female*_ = 28 vs. *n*_*male*_ = 9). To control this effect, we re-analyzed the Face-to-Art condition by equalizing the groups and using the full participant pool as a proxy for aesthetic preference (Fig. 4). Under these conditions, the average MSE was 0.88 (±0.27) for males, 0.94 (±0.61) for females. The average correlation coefficient was *r* = 0.29 ± 0.20 for males, *r* = 0.35 ± 0.22 for females. These results indicate that the observed performance gap between genders was less notable when sample sizes were normalized.

## 5 DISCUSSION

### 5.1 Consistency of aesthetic preferences

In this study, we demonstrated the consistency of aesthetic preferences across multiple domains, extending the contemporary Theory of Aesthetic Preferences (Miller and Hübner (2020); Miller et al. (2025)). We introduced a novel approach in which predicting individual outcomes based on aesthetic preferences is framed as a typical collaborative filtering task. This framework provides a computationally verifiable method for testing aesthetic preference across diverse domains. This framework can be integrated with advanced modeling techniques, such as aesthetic neural networks (Liu et al. (2019)), reinforcement learning frameworks (Aleem et al. (2020)), deep convolutional networks (Iigaya et al. (2023)), and a variational autoencoder (VAE) with meta-learning (Chen and Gong (2025)), to further enhance predictive performance.

Our behavioral analysis revealed high correlations between the “natural” domain (male/female faces and scenes), whereas correlations were notably lower between these domains and artworks (Fig. 3C). These findings align with Vessel et al. (2018), who also demonstrated the distinction between “natural” (faces, scenes) and “artificial” (artworks) domains. While the cross-domain test correlations using our proposed approach were relatively high (Fig. 4, 5), they did not reach the levels observed in within-domain tests. This discrepancy suggests the presence of the domain-specific modulations that were not accounted for in the current model. Consequently, the quantitative evaluation of the tension between domain-specific and domain-universal modulations remains a critical step for applying aesthetic preference models to real-world challenges, such as recommendation systems Liu et al. (2019).

Prior research has provided robust evidence that the aesthetic preferences differ with gender within a domain Miller and Hübner (2022). In this study, we examined whether gender also modulates cross-domain performance. Unexpectedly, we observed a modest increase in MSE and a concomitant decrease in test correlation for the male group in art prediction cases ((Fig. 6A, 7A)). However, in scene prediction, the male group demonstrated a significant increase in MSE and a decrease in test correlation when using their responses to Female faces as a proxy for aesthetic preferences. The observed disparity between art and scene predictions likely stems from the closer ecological or visual relationship between scenes and faces compared to artworks (Vessel et al. (2018)). However, we emphasize that our current male sample was limited; consequently, these results should be interpreted as preliminary evidence of gender-specific modulation rather than a definitive conclusion.

We posit that the male-specific modulation identified here may be attributed to a “gender attractiveness gap”. Recent evidence from Wassiliwizky et al. (2025) suggests that female faces are typically rated higher by both males and females; such a behavioral trait might systematically inflate the male participant’s responses beyond their intrinsic aesthetic preferences. To rigorously test this hypothesis, future research should incorporate explicit attractiveness metrics to investigate the relationship between them and the preference scores.

### 5.2 Neural basis of cross-domain aesthetic preferences

To further bridge our behavioral findings with established neurophysiological models, we propose a simplified circuit for cross-domain aesthetic evaluation (Fig. 8). In this study, the evaluative process (ranking/rating) occurred post-stimulation, a temporal window typically associated with the functional recruitment of the OFC and broader PFC (Ticini (2017)).

**Figure 8.**
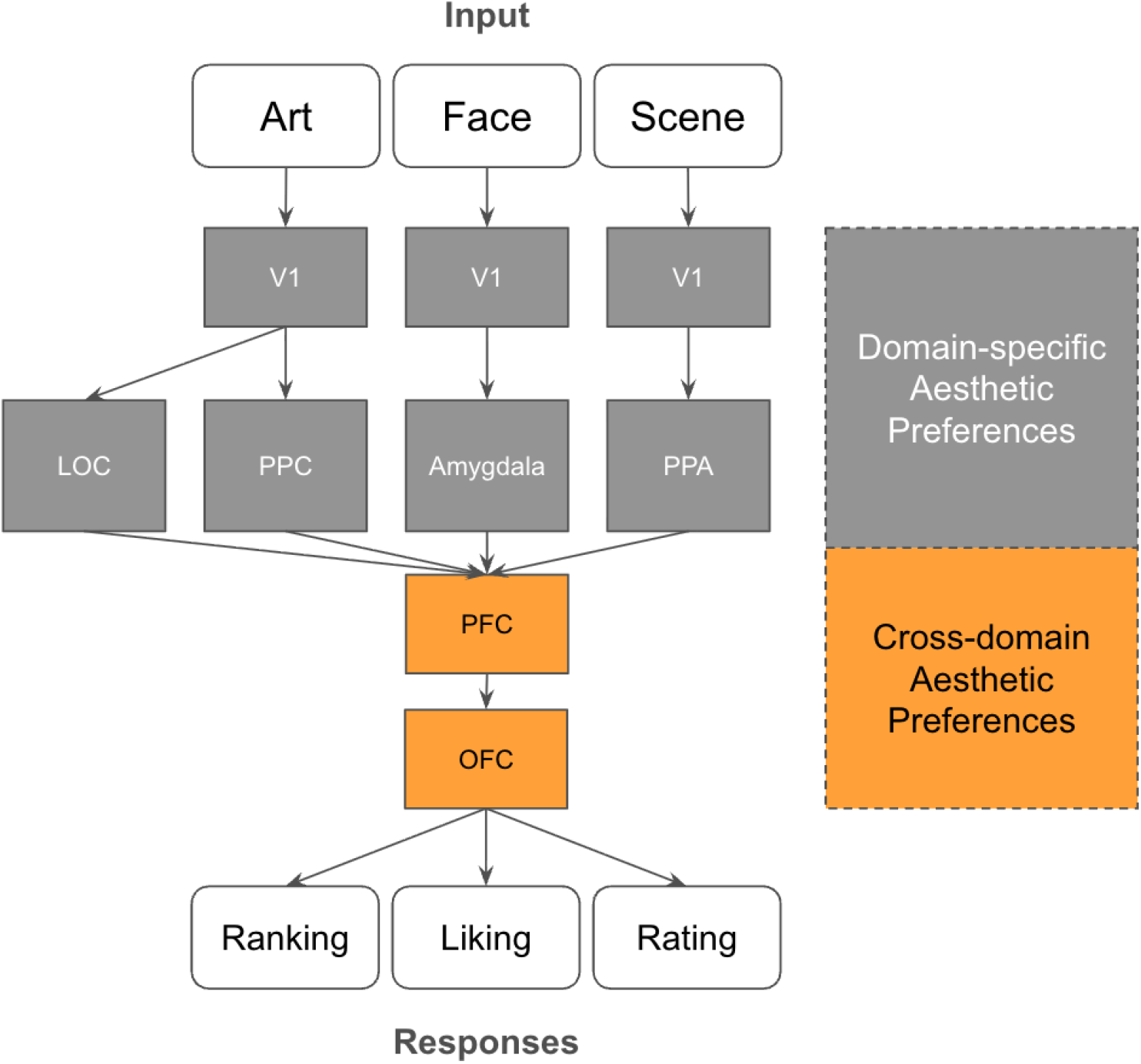
Functional circuit model of cross-domain aesthetic processing. The diagram illustrates the convergence of domain-specific pathways (LOC/PPC for Art, Amygdala for Faces, and PPA for Scenes) onto centralized valuation hubs in the PFC and OFC. The transition from grey to orange blocks represents our hypothesis that stable, cross-domain aesthetic signatures emerge through the integration of these disparate sensory inputs into a domain-invariant value signal.

The pathways depicted in Fig. 8 illustrate the transition from domain-specific sensory processing to centralized value integration. For instance, artworks are processed through the primary visual cortex (V1) and evolve toward the OFC via the lateral occipital cortex (LOC) and PPC (Iigaya et al. (2023)). In contrast, facial aesthetic judgments rely heavily on the amygdala (Adolphs et al. (1998); Adolphs and Tranel (1999)), with established functional connectivity to the OFC (Todorov et al. (2013)), while scene evaluation is bridged via the parahippocampal place area (PPA; King et al. (2002); Sewards (2011)).

Our behavioral evidence for cross-domain consistency implies that while early-stage ‘upper’ pathways may maintain domain-specific aesthetic preferences, these signals converge into a stable, domain-invariant representation within the PFC and OFC. This architecture suggests that these prefrontal regions act as a functional hub for the universal encoding of aesthetic value. However, the exact extent to which these underlying neural representations remain truly invariant across modalities remains a critical question for future neuroimaging research.

The specific neural pathway of aesthetic evaluation highly resembles the subjective value hierarchy described by Pham et al. (2021). This valuation axis emerges from primary sensory areas and progresses towards the terminals of the Default Mode Networks (DMN), following the established principal cortical gradient of macroscale organization Margulies et al. (2016). Taken together, these findings suggest a potential role of DMN in mediating the cross-domain aesthetic preferences. This hypothesis is supported by the work of Vessel et al. (2012, 2013) who found that the intense aesthetic experiences in the art domain were found associated with a distinct lack of deactivation in DMN. This suggests that the DMN may serve as a site where external stimuli are integrated with internal, stable representations of personal preference.

Future studies should consider using high-resolution neuroimaging and electrophysiological techniques, such as fMRI and electroencephalography (EEG), in combination with explainable artificial intelligence (XAI). Such an approach would allow for a deeper investigation into whether the latent representations of aesthetic preference in the human brain remain consistent across diverse stimulus categories, ultimately confirming the existence of a domain-invariant aesthetic ‘signature’ in the human brain.

### 5.3 Limitations

Despite the robust predictive performance of our framework, several limitations warrant consideration. First, the domains evaluated in this study were exclusively within the visual modality. While our results demonstrate cross-domain consistency within vision, further investigation is required to determine whether these aesthetic principles extend to other modalities, such as the auditory or haptic (touch) domains. Testing for cross-modal consistency would provide a more comprehensive understanding of whether aesthetic preference constitutes a truly centralized, modality-independent trait.

Second, the demographic homogeneity of our sample may limit the generalizability of these findings. All participants were of Japanese nationality and shared a similar cultural background. Given that cultural exposure significantly shapes aesthetic standards (Masuda et al. (2008)), the degree of cross-domain consistency observed in this study warrants validation across diverse cultural cohorts to confirm its broader universality. Furthermore, our sample was predominantly female, which precludes a comprehensive analysis of potential gender-based differences in these behaviors. The findings related to gender must be considered exploratory. Future research should utilize larger, more balanced datasets to reach robust conclusions regarding the extent to which gender modulates cross-domain aesthetic consistency.

Finally, the experimental design utilized isolated tasks, which lacked real-time social interaction. This limits the interpretation of our findings within the context of dynamic social environments. In real-world social contexts, the “attractiveness halo effect”, whereby attractive individuals are implicitly associated with positive personality traits, is a pervasive bias that significantly modulates behavioral outcomes (Zaki et al. (2011)). It remains unknown how the presence of others or the explicit awareness of peer responses might trigger mechanisms of social contagion and conformity, potentially overriding the cross-domain consistency observed in this study.

Future research should incorporate social-influence paradigms or “hyper-scanning” approaches to explore how individual preference consistency interacts with collective social pressures. Investigating whether these external social cues recruit distinct or overlapping neural circuits within the reward and social-cognition networks will be essential for a comprehensive model of aesthetic-driven social behavior.

## 6 CONCLUSION

In this study, we investigated the cross-domain consistency of aesthetic preferences through a novel computational lens. By framing aesthetic evaluation as a collaborative filtering task and leveraging inter-subject similarity matrices, we demonstrated that an individual’s aesthetic judgments are not merely isolated responses to specific stimuli. Instead, they reflect a stable, latent trait that allows for the accurate prediction of behavior across disparate domains, including art, faces, and scenes.

Our findings provide empirical support for the TAP and extend the concept of aesthetic universality from a cross-cultural context to a cross-domain framework. The high predictive accuracy of our model, even when transitioning between “natural” domains like faces and “artificial” domains like artworks, suggests that the human brain encodes aesthetic value as an abstract, domain-independent signal. This conclusion is further supported by current neurophysiological models of the OFC and the DMN, which function as high-level integration hubs for subjective value.

While gender-specific modulations and the limitations of a single-culture sample suggest areas for future refinement, the core results underscore the robustness of aesthetic profiles as a component of social identity. Beyond theoretical contributions, this research offers a practical foundation for the next generation of personalized recommendation systems and social behavior analytics. By capturing the “latent signature” of a user’s taste in one modality, digital platforms can more effectively personalize experiences across entirely different domains, ultimately bridging the gap between computational modeling and the complex reality of human aesthetic experience.

## AUTHOR CONTRIBUTIONS

Conceptualization: T.Q.P., J.C. Methodology: T.Q.P., J.C. Investigation: T.Q.P. Writing - Original Draft: T.Q.P. Writing - Review and Editing: T.Q.P., J.C. Supervision and Funding Acquisition: J.C. All authors reviewed the manuscript.

## DATA AVAILABILITY STATEMENT

Due to copyright and privacy constraints, the stimulus set, generated data, and codes are available from the corresponding authors upon reasonable request for research purposes only.

